# Challenges in estimating the motility parametersn of single processive motor proteins

**DOI:** 10.1101/148346

**Authors:** F. Ruhnow, L. Kloß, S. Diez

## Abstract

Cytoskeletal motor proteins are essential to the function of a wide range of intracellular mechanosystems. The biophysical characterization of their movement along their filamentous tracks is therefore of large importance. Towards this end, single-molecule, *in vitro* stepping-motility assays are commonly used to determine motor velocity and run length. However, comparing results from such experiments has proved difficult due to influences from variations in the experimental conditions and the data analysis methods. Here, we investigate the movement of fluorescently-labeled, processive, dimeric motor proteins and propose a unified algorithm to correct the measurements for finite filament length as well as photobleaching. Particular emphasis is put on estimating the statistical errors associated with the proposed evaluation method as knowledge of these values is crucial when comparing measurements from different experiments. Testing our approach with simulated and experimental data from GFP-labeled kinesin-1 motors stepping along immobilized microtubules, we show (i) that velocity distributions should be fitted by a *t* location-scale probability density function rather than by a norm^*^al distribution, (ii) that the impossibility to measure events shorter than the image acquisition time needs to be accounted for, (iii) that the interaction time and run length of the motors can be estimated independent of the filament length distribution, and (iv) that the dimeric nature of the motors needs to be considered when correcting for photobleaching. Moreover, our analysis reveals that controlling the temperature during the experiments with a precision below 1 K is of importance. We believe, our method will not only improve the evaluation of experimental data, but will also allow for better statistical comparisons between different populations of motor proteins (e.g. with distinct mutations or linked to different cargos) and filaments (e.g. in distinct nucleotide states or with different posttranslational modifications).

## Introduction

Cytoskeletal motor proteins are essential for long-range intracellular transport [1], the malfunction of which can cause a number of pathologies including neurodegenerative diseases [2]. The precise characterization of motor proteins with regard to their intrinsic function and the investigation of factors that influence their behavior thus constitutes an important part of medical and biophysical research. To study motor proteins in minimal *in vitro* systems, fluorescence imaging of single-motors, stepping along their filamentous tracks, still remains at the forefront of biophysical tools [3-6]. However, quantitative estimation of the crucial motility parameters, namely velocity, interaction time and run length, proves to be challenging owing to fundamental limitations in the experimental design. Because the processive run of a motor may be prematurely terminated by the end of a filament or the motor may be rendered invisible due to photobleaching of the attached fluorescent marker, so-called censored events will be part of any experimental data [7]. As these censored events are prone to significantly bias the results, reliable correction methods are needed. Although corrections for finite filament length and photobleaching have been investigated individually in the past [8-11] the field still lacks a unified methodology. Here, we suggest an approach, which addresses the above-mentioned challenges. Besides evaluating experimental data obtained from the motility of single, GFP-labeled kinesin-1 motors, we perform numerical simulations using a priori known parameters to show how statistical analysis allows for characterizing the certainty of a given measurement. Knowledge of the latter is of crucial importance when data from different measurements are to be compared. Along with our findings, we emphasize the importance of a precise temperature control during the measurements and describe a detailed workflow, which facilitates the analysis of experimental data.

## Materials and Methods

### Motor proteins and filaments

Histidine-tagged, truncated (1-430 a.a.) rat kinesin-1 labeled with eGFP (rKin430-eGFP) was expressed and purified as previously described [12]. Porcine tubulin was purified from porcine brain (Vorwerk Podemus, Dresden, Germany) using established protocols [13]. Microtubules were grown for 2 hours at 37°C from a 80 μl BRB80 (80 mM Pipes [Sigma], pH 6.9 adjusted with KOH [Merck], 1 mM EGTA [Sigma], 1 mM MgCl_2_ [Merck]) solution supplemented by 2 mM tubulin (75% unlabeled and 25% rhodamine labeled [TAMRA; Thermo Fisher Scientific]), 1mM GMP-CPP (Jena Bioscience, Jena, Germany) and 1mM MgCl_2_. 80 μl of the microtubule solution was centrifuged in a Beckman Airfuge (A95 rotor; Beckman, Brea, CA) at 100000*g* for 5 min. The pellet was resuspended in a volume of 200 μl BRB80T (BRB80 supplemented by 1 mM taxol [Sigma]). The solution was kept at room temperature over night and the 200 μl microtubule solution was centrifuged and resuspended (in 200 μl BRB80T) again before the experiment.

### Single molecules stepping assay

The employed stepping assays using TIRF microscopy have been extensively described by Korten et al. [14]. Briefly, we performed the experiments in flow channels [14], self-built from two glass coverslips (22x22mm^2^ and 18x18mm^2^; Corning, Inc., Corning, NY), which were cleaned in piranha solution (H_2_O_2_/H_2_SO_2_, 3:5; both purchased from Sigma), silanized with 0.05% dichlorodimethylsilane in trichloroethylene (Sigma) and glued together by heated pieces of Parafilm M (Pechiney Plastic Packaging, Chicago, IL). The flow sequence was as follows: (i) The flow channel was filled with a solution of TetraSpeck microspheres (diameter 100 nm; Thermo Fisher Scientific) diluted 200-fold in BRB80, which were used for drift and color correction [16]. (ii) After 2 min, the solution was exchanged with a BRB80 solution containing 77.5 μg/ml anti-*β*-tubulin antibodies (SAP4G5; Sigma). (iii) After 5 min, the surface was blocked with a solution with 1% Pluronic F-127 (Sigma) in BRB80 for 15 min. (iv) Microtubules diluted 10-fold to prevent microtubule intersections, were incubated for 5 min to bind to the tubulin antibodies. (v) The microtubule solution was finally replaced by the motility solution (BRB80 containing 10 μM taxol, 0.04mM glucose [Sigma], 0.2mg/ml glucose oxidase [SERVA],0.02 mg/ml catalase [Sigma], 10 mM DTT [Fermentas], 0.1mg/ml casein [Sigma], 10mM Mg-ATP [Sigma]) supplemented by 4 μg/ml rKin430-eGFP. For bleaching time estimation, undiluted dimly labeled microtubules were used (10x less rhodamine labeling) and ATP was replaced with 10 mM AMP-PNP (Sigma) in all solutions.

### Optical Imaging

Fluorescence imaging was performed using an inverted fluorescence microscope (Zeiss Observer Z1; Zeiss, Jena, Germany) with a 100x oil immersion objective (Zeiss APOCHROMAT; numerical aperture 1.46; Zeiss) and an additional 1.33x magnifying optovar. The final pixel size was 117 nm. Microtubules were observed by epifluorescence using a Lumen 200 metal arc lamp (Prior Scientific Instruments Ltd., Fulbourn, UK) with a TRITC (exc 534/30, em 593/40, dc R561; all Chroma Technology, Rockingham, VT) filter set. rKin430- eGFP motor proteins were observed in total internal reflection fluorescence (TIRF) mode by using a PhoxX 488 nm Laser (Omicron-Laserage, Rodgau-Dudenhofen, Germany) with a GFP (exc 470/40, em 525/50, dc 495; all Zeiss) filter set. Image acquisition was performed at 100 ms exposure time in streaming mode by an electron-multiplied charge-coupled device camera (iXon Ultra DU-897U; Andor, Belfast, Northern Ireland) in conjunction with a Metamorph imaging system (Universal Imaging Corp., Downingtown, PA). The temperature was measured with a temperature sensor (IT-23; Physitemp Instruments, Inc., Clifton, NJ) directly in the flow channel, which was connected to multipurpose thermometer (BAT-10; Physitemp Instruments, Inc.). Temperature control was implemented using a custom-made hollow brass ring (MPI-CBG Mechanical Workshop, Dresden, Germany) around the objective connected to a water bath with combined cooling and heating unit (F25-MC Refrigerated/Heating Circulator; JULABO GmbH, Seelbach, Germany). While the electric components of the microscope setup usually increase the room temperature as well as the temperature of the microscope body, our temperature control kept the temperature in the flow channel stable within 0.5 K over hours. See Supporting Material S1 for further experimental considerations.

### Single Molecule Analysis

Single kinesin-1 molecules and microtubules were tracked using FIESTA [17]. Molecules that showed any pauses or stalling were disregarded. After drift and color offset correction, the molecule position was projected on the microtubule centerline (see Supporting Material S2 for detailed instructions on tracking with FIESTA). The resulting distance along the centerline as well as the detachment position was utilized for further evaluation in different MATLAB (Mathworks, Natick, MA) scripts. For velocity simulations a Monte-Carlo-Simulation of a Poisson Stepper was used and to create exponential distributions the MATLAB function *exprnd* was employed. The evaluation of the cumulative distribution function utilized *ecdf*, which also includes the optional Kaplan-Meier-Estimator, and for least-square fitting fit was used. Bootstrapping was done with *parfor* included in the *Parallel Computing Toolbox* of MATLAB. The MATLAB code for the simulations as well as for the evaluation can be found in Supporting Material S3 & S4.

### Bootstrapping Method

The statistical error of an evaluation method with a limited number of measurements can be described using a bootstrapping method [4]. Briefly, from the data set individual measurements are randomly selected with replacement. Here, the complete data set is always available when picking the measurement, which means that any measurement can also be selected more than once. Now, the new randomly selected data set is analyzed using the desired evaluation method. This procedure is repeated for sufficient number of repetitions (e.g. n = 100) with randomly selected data sets. The resulting bootstrapping distribution can be described by a normal distribution with the mean denoting the actual result and its standard deviation describing the statistical error. This statistical error is only the result from random sampling and describes the error that is to be expected when repeating the experiment.

## Single Motor Protein Stepping Assay

Motor proteins moving along their filaments can be described theoretically as Poisson steppers (Figure 1). A simplified model of the motor protein kinesin-1 stepping along microtubules (MTs) is shown in Figure 1A. While the attachment rate 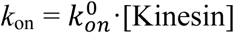 is influenced by the motor protein concentration [Kinesin] in solution, the detachment rate *k*_off_ only depends on the motor-filament interaction. This interaction is described by (i) the motor velocity *v*, which can be used to derive the stepping-rate *k*_step_ = *v*/*d* with *d* denoting the step size, (ii) the interaction time *τ*, which can be used to derive the detachment rate *k*_off_ = 1/*τ,* and (iii) the run length *R,* which can be used to link the detachment rate *k*_off_ to the mechano-chemical cycle of the motor protein (e.g. *R* = *v* · *τ*). The first challenge in the determination of these parameters can be seen in the experimental kymograph in Figure 1B. There, clear linear motion can be observed for some motors (e.g. red box), while it is unclear if short interactions are actual movement or unspecific interactions (e.g. blue box). In our analysis we therefore require a motor to be visible for five or more consecutive imaging frames and to move over a distance longer than the size of two pixels without pausing. While these experimenter-defined thresholds appear arbitrary, we will show that their choice does not affect the results.

**Figure 1:**
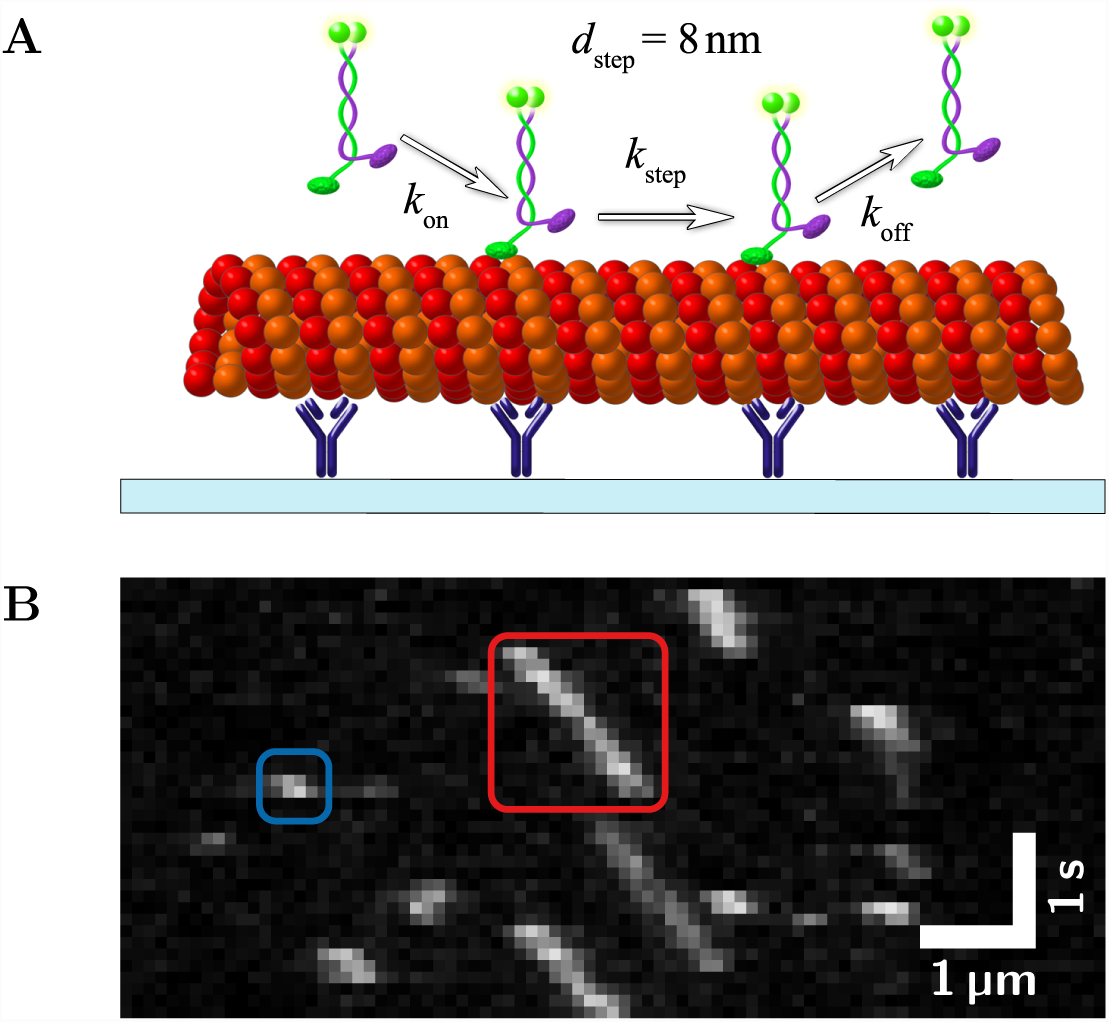
Single-motor stepping assay: (A) Schematic depiction of stepping assay. GFP-labeled kinesin-1 motors move along rhodamine-labeled microtubules that are immobilized by antitubulin antibodies to a glass surface. (B) Typical kymograph (experimental data) of a stepping assay with kinesin-1. The red box shows a single motor that is measurable (interaction time *τ* = 1.3 s), the blue box shows a motor with an interaction time that is too short for a reliable measurement. In the latter case it is unclear whether the motor moved processivly along the microtubule or interacted only unspecifically.

In the following we will describe the procedures to estimate velocity, interaction time and run length. While estimating the mean velocity will be rather straightforward, we will show that it is more challenging (but equally important) to estimate the associated statistical error. As motor velocity is a good reporter of the environmental conditions (foremost temperature but also ionic strength and pH) knowledge about both, mean velocity and error, is of importance when potentially combining data from different fields of view or different experiments. Without taking precautions, the environmental parameters can vary significantly over the course of an experiment and unwanted influences can be minimized by pooling data from measurements with non-differing velocities only. For reducing the systematic error in estimating the interaction time and run length, data analysis needs to address additional experimental challenges such as limited filament length and photobleaching.

## Evaluation of the velocity

The velocity of individual processive motors is determined by tracking their positions over time, *X(t)* and *Y(t)*. These positions are projected on the centerline of the filament and the distance *D(t)* the motor moved along the filament is calculated. Fitting *D(t)* with a linear function *D(t)* = *v*·*t*+*c* yields the velocity *v*. A typical experimental velocity distribution of *N* = 543 kinesin-1 motors is shown in Figure 2A. Although, upon first sight the distribution resembles a normal distribution, hypothesis testing reveals that the data can not be described by a normal distribution (*p* < 0.001, Kolmogorov-Smirnov test).

**Figure 2:**
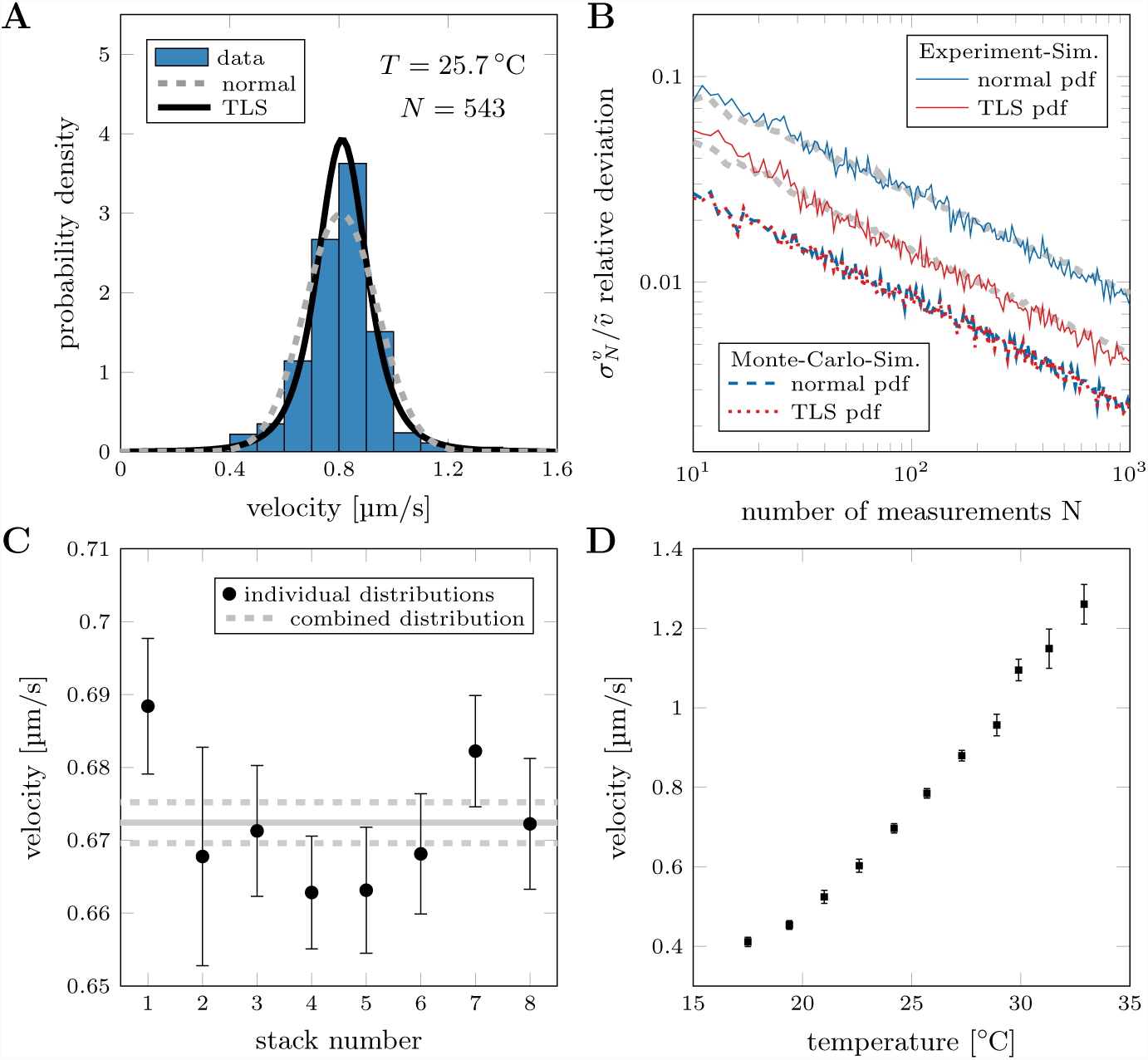
Evaluation of velocity in single-motor stepping assays: (A) Histogram of kinesin-1’s velocity distribution at temperature *T* = 25.7 °C with corresponding normal and TLS pdf (estimated with MLE). (B) Evaluation of simulations using normal and TLS pdf. The red/blue dashed lines show the results using the Monte-Carlo simulation whereas the solid lines depict results from simulations that resemble more realistic experimental conditions (Monte-Carlo simulation with spatial averaging and additional positional error). Whereas both distributions yield the same precision for the results obtained from the Monte-Carlo-Simulation, the TLS pdf is more precise than the normal pdf with simulated experimental data. The gray dashed lines are the average bootstrapping errors (see Methods), used to estimate the statistical errors. (C) Velocity during acquisition of one data set in 8 different fields of view. The temperature was kept constant within 0.5 K (23.5 - 24 °C) and the velocity shows only marginal deviations over time. Therefore, data can be pooled to create a combined data set (grey lines, dashed lines indicate error, *N*_total_ = 5208). (D) Dependence of velocity on temperature (measured in the flow channel). A temperature increase of 1 K increases the velocity by more than 5 %.

To investigate the reason behind this discrepancy, we created a Monte-Carlo-Simulation with 10000 traces of motor proteins stochastically stepping with a rate of *k*_step_ = 100 s^−1^ and step size *d*_step=_ 8 nm (*k*_off_ = 0, total time per trace 20 s). We looked at the number of steps *N*_steps_ taken by each of these simulated motor proteins at specific time points (e.g. *t*_1_ = 1 s, *t*_2_ = 3 s, etc). At each time point, *N*_steps_ is described by a Poisson distribution, which can be approximated with normal distributions (*N*_steps_ > 10; Figure S1A). Since the velocity is described by *v* = *N*_steps_ · *d*_step_ /*T* the mean velocities are the same at each time point, but the widths of the normal distributions vary (Figure S1B).

If the detachment rate is changed to *k*_off_ = 0.5 s^-1^, each motor has a different interaction time and thereby the velocity distribution of all motors is a mix of normal distributions with the same mean values but different widths. In general, motors with shorter interaction times have a higher variance in the velocity distribution than motors that interact longer (under the same imaging conditions, Figure S1B). Consequently, the observed velocity distribution is not a normal distribution.

To further adjust the simulation to more realistic experimental conditions, spatial averaging over the positions *D(t)* during the acquisition time of individual imaging frames (e.g. 100 ms accounting for a finite frame rate of *f* = 10 s^-1^) was performed and a positional error (due to tracking uncertainty) was incorporated by adding normally distributed noise (*σ* = 100 nm). Due to the finite acquisition time, no exact information about the attachment and detachment time can be extracted. Consequently, the positional averaging during acquisition of these frames would bias the estimation towards slower velocities (see Supporting Material S5). Therefore, the first and last tracked frames have to be excluded in the linear regression of *D(t)*.

After obtaining a single velocity for each motor, a mean velocity *v* for one experiment can be obtained by estimating the characteristic parameters of a probability density function (pdf) with Maximum-Likelihood-Estimation (MLE). As pdf we used a *t* location-scale (TLS) distribution [18], which includes a shape parameter v in addition to the location *μ* and scale *σ* parameters, which are conventionally used to describe a normal distribution. The TLS pdf that MATLAB uses in MLE is described by the following equation:

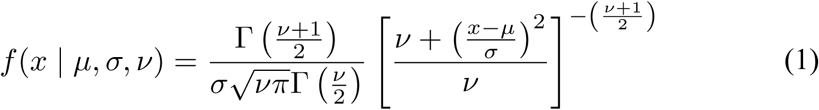

As can been seen in Figure 2A, the TLS pdf fits the velocity distribution better than the normal distribution because it accounts for heavier tails (see also Figure S2A). In order to compare the statistical errors when using these pdfs, we picked out a random set of *N* motor proteins from our simulation (with replacement, see bootstrapping in Methods). We calculated the velocity for each motor, and estimated the mean velocity 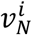 for that random set using both normal and TLS pdfs. This procedure was repeated *n* = 100 times and the deviation from the true velocity 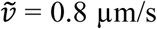 (*k*_step_ = 100 s^-1^ and *d*_step_ = 8 nm) was calculated:

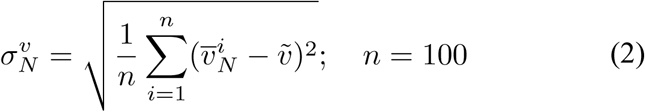

The relative deviation 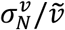 is plotted as function of the number of measurements in Figure 2B. Although there is only a small difference between using a normal or TLS pdf when evaluating the Monte-Carlo-Simulation, this difference increases when evaluating time-averaged traces with additional positional error (corresponding to experimental data). It turns out that, to reach a relative error Δ*v/v* of 2% (with 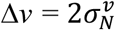, confidence interval 95%) only 200 measurements are necessary when using a TLS pdf compared to 750 measurements when using a normal pdf. Here, the difference between TLS pdf and normal pdf depends on the average number of data points in each trace available (defined by time resolution and interaction time) for each trace as well as on the positional error. On average, using either distribution yields the same mean velocity, but the TLS pdf yields a smaller relative error because it fits the velocity distribution significantly better [ 18] (see Supporting Material S5 for a more extensive investigation of the velocity estimation).

#### Workflow for estimating velocity

I. Extract the distance vs. time information for each motor protein (see Supporting Material S2 for instructions on how to track and analyze single fluorescent motor proteins using FIESTA [17]).
II. Fit distance vs. time trace with linear regression leaving out first and last frame of the trace.
III. Use MLE to fit a TLS pdf to the velocity distribution.

To calculate the error of the mean velocity *v* with the TLS pdf either use the 95% confidence interval (estimated with MLE) or bootstrapping (see Methods). See MATLAB code in the Supporting Material S4.

In addition to measuring the velocity precisely it is also essential to limit any external influence on the motility parameters, most importantly the temperature. Conventionally, velocity measurements on motor proteins are performed at ‘room temperature’ and the actual temperature is not specified precisely. However, temperature variations can lead to small but significant velocity differences within one experiment and only by actively stabilizing the temperature in the flow channel using a temperature control system (within ±0.5 K, see Supporting Material S6) we could remove the temperature influence on the motor stepping (Figure 2C). In contrast, not accounting for temperature differences when comparing data sets (e.g. from different days or labs) can lead to a grave misinterpretation of data. Even a 1 K temperature increase can influence the velocity by more than 5 % (Figure 2D). The temperature effect becomes even more prominent for measurements at ‘room temperature’ (e.g. data points at 24.2 to 25.7°C) with about 13 % increase in velocity over 1.5 K. Thus, measuring and reporting both temperature and velocity is utmost importance when comparing the motility parameters of motor proteins. In turn, the motor velocity can be used as a control parameter to assure a constant temperature. In fact, together with our temperature control system this strategy enabled us to pool experimental data sets from different image sequences and fields of view into a combined data set for further analysis.

## Evaluation of the interaction time and run length

Interaction time and run length of motor proteins are theoretically exponentially distributed [19] and our Monte-Carlo-Simulations indeed show exponential distributions for both parameters. This can be attributed to a stochastic detachment with rate *k*_off_ where the interaction time is *τ* = *k*_off_^-1^ and the run length is *R* = *v* · *k*_off_^-1^. In order to test different methods to evaluate censored data, we simulated exponential distributions and evaluated them using three different methods previously described in the literature: (i) Least-squares-fitting of the probability density function (LSF-PDF) where the data is binned in a histogram, and the locations and heights of the bins are fitted by an exponential pdf 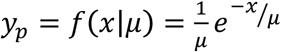 [19]. (ii) Least-squares-fitting of the cumulative distribution function (LSF-CDF) where the data is used to create a cumulative probability distribution that is fitted with the cumulative distribution function (cdf) 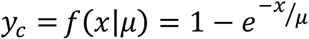 [4]. (iii) Maximum-Likelihood-Estimation (MLE) where the data is used directly to estimate the parameters of the exponential distribution. For complete exponential distributions (where all events are measurable) the MLE method yields the most precise results, because it can be solved analytically (for exponential distributions among others). In contrast, least-squares-fitting involves numerical optimization of the parameters, which is terminated when a certain tolerance is reached. This slightly decreases the precision with which the parameters are estimated. However, the experimental limitations prevent us from measuring a complete exponential distribution. On one hand, it is impossible to include motility events with *τ* shorter than the time resolution (in our case 100 ms). And on the other hand, events with short interaction times might be easily discarded as noise during the evaluation procedure. Due to the missing short events, the measured exponential distribution is not complete and the evaluation method has to be adjusted. Using LSF-PDF, the first bin is underrepresented and needs to be disregarded when fitting the pdf (Figure 3A). Using LSF-CDF, a cutoff parameter *x_0_* has to be introduced in the cdf 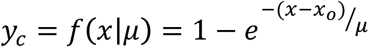 [4] to account for the missing measurements (Figure 3B). The cutoff parameter can be set as a constant (LSF-CDF(static), [4]) or as a free fit parameter (LSF-CDF(free), [5]). In MLE it is not possible to correct for missing events. Here, we characterize the evaluation of exponential distributions with LSF-PDF, LSF-CDF and MLE by creating a random data set of an exponential distribution with 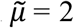 (we disregard the difference between run length and interaction time for now because the following applies to any exponentially distributed data). Analogous to the velocity in the previous section we analyze the deviation of the estimated mean 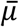 (from *n* = 100 independent data sets) from the true 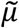 in dependence of the number of measurements *N*:

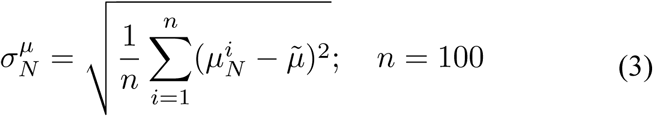

**Figure 3:**
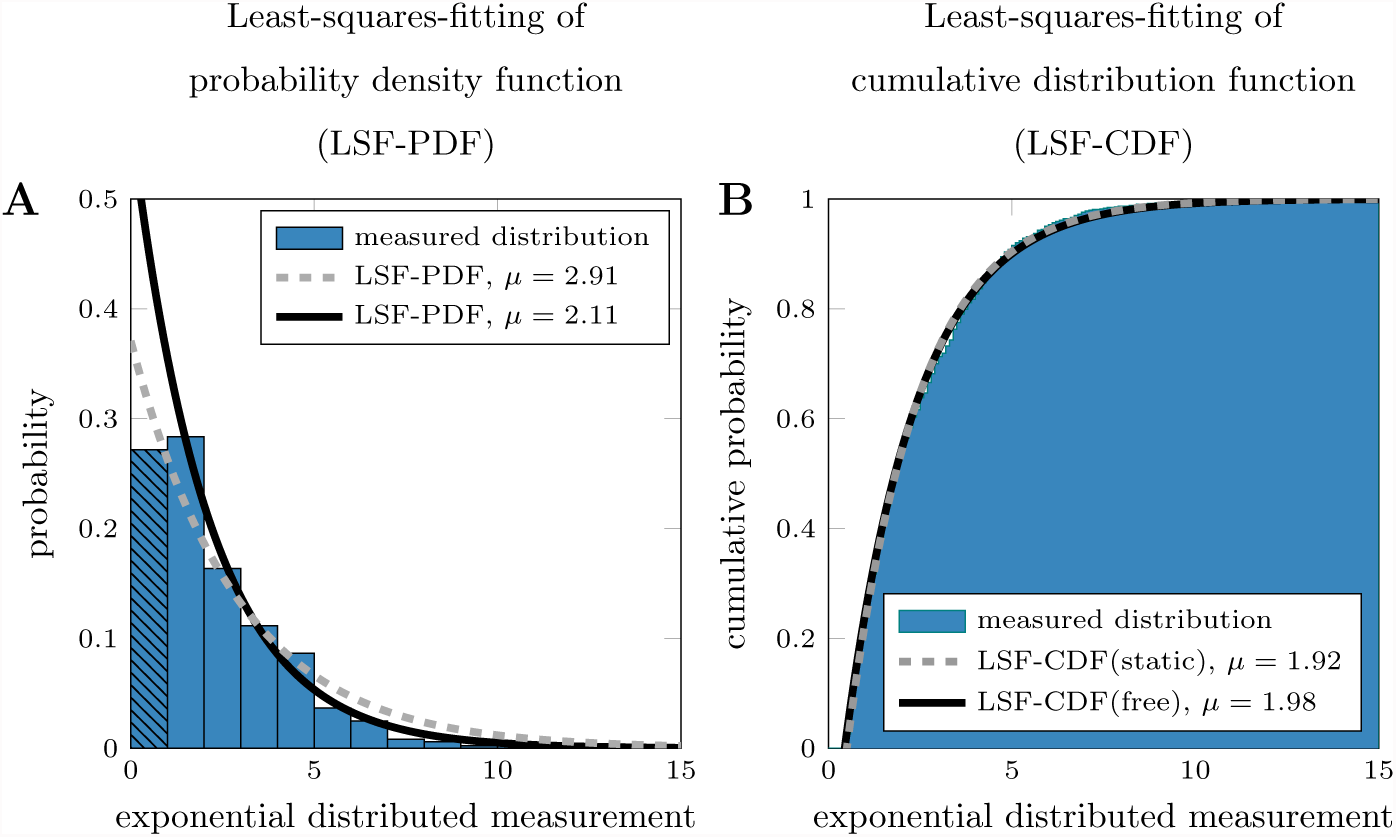
Modified exponential distribution to simulate real measurements: (A) Modified exponential probability distribution with *μ* = 2 and *N* = 1000. Here, the first bar is underrepresented (black pattern), due to experimental limitations. The dashed line shows the LSF-PDF of the complete distribution, whereas the solid line shows the fit with the first bar excluded. (B) Modified cumulative probability of an exponential distribution with *μ* = 2 and *N* = 1000. Here, all measurements *x* < 0.5 are disregarded, due to experimental limitations. The grey dashed line shows the LSF-CDF(static) of the distribution with a fixed *x_0_*, whereas the solid black line shows the fit with *x_0_* as free fit parameter (LSF-CDF(free)).

Figure 4A shows the results when using the complete exponential distribution. The relative deviation depends only on the number of measurements due to the statistical nature of the exponential function. As expected, all methods show a clear 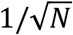 behavior, but the LSF-PDF yields a higher statistical error due to ‘coarse graining’ by binning. Here, the data is binned, but the distribution of the measurements within the bins is skewed (because an exponential distribution is continuously decreasing) which results in a deviation from the expected result. If the simulation is adjusted to represent realistic experimental results (Monte-Carlo simulation with spatial averaging, additional positional error and missing short events) two methods do not show this characteristic behavior (see Figure 4B). Here, MLE has the highest systematic error because sampling of the whole distribution is necessary and also using LSF-CDF(static) can yield a systematic error. Because the cutoff is chosen arbitrarily, it would force the cdf through a fixed point on the x-axis. In experiments, the exact cutoff could be hidden within the time resolution or tracking accuracy. Therefore, it is essential to use the cutoff *x_0_* as a free fit parameter when evaluating the exponential function. The LSF-PDF is again influenced by ‘coarse graining’ and therefore yields a higher statistical error, which leaves the LSF-CDF(free) as the most precise method for evaluation, without any systematic error that could bias the result. Note that the statistical error can not be smaller than 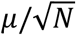, even if the confidence interval of the fitting is smaller. The fitting error from least-squares-fitting occurs in addition to the statistical error, which results from the random sampling of the exponential distribution. Hence, it is more important to measure a large number of data points rather than improving the precision of the individual measurements. The code for fitting a particular experimental data set can be found in the Supporting Material S4 along with an extensive discussion on the creation of the cdf and least-squares-fitting in Supporting Material S7. In the following, we only use the LSF-CDF(free) to evaluate exponential distributions.

**Figure 4:**
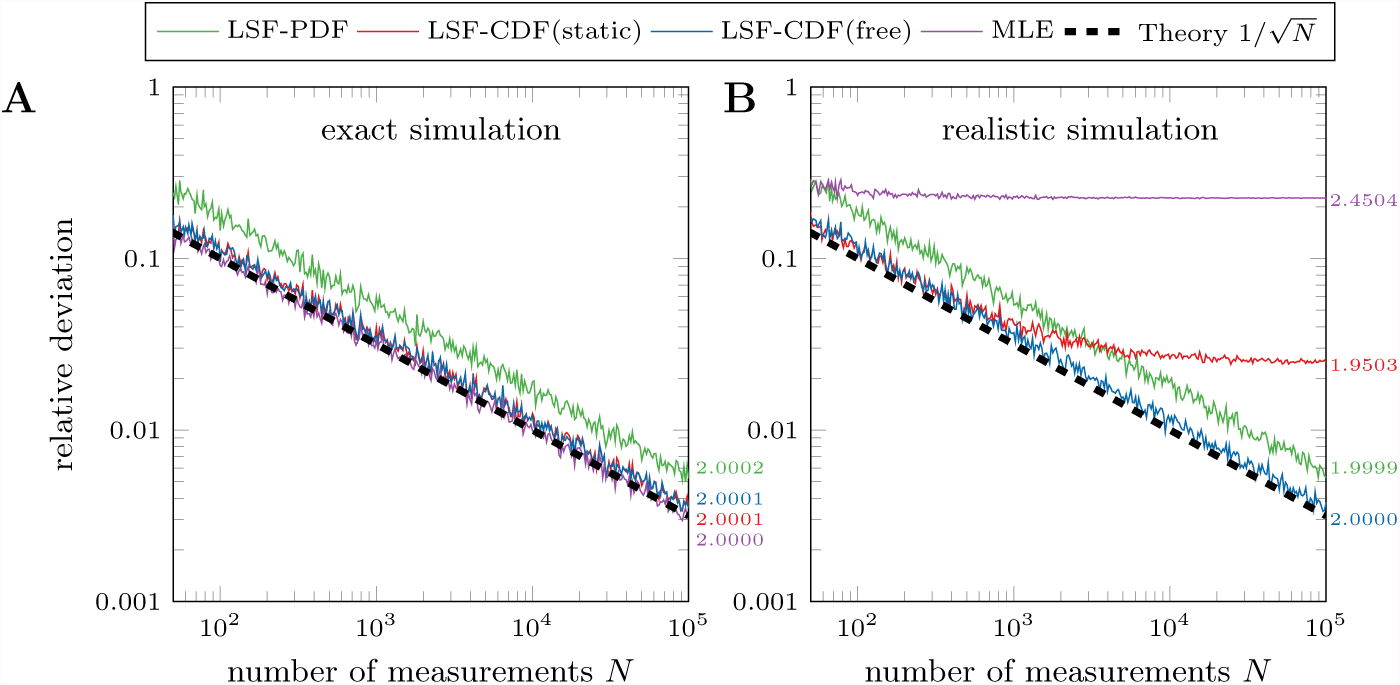
Comparison of methods to estimate the parameters of exponential distributions: (A) Relative deviation from the a priori value of the simulation estimated using an exact and complete exponential distribution. All methods reach the right result (see values on the right), only the LSF-PDF method has a higher statistical error due to ’coarse graining’ by binning. (B) Same relative deviation, but using a modified exponential distribution with a resolution of 0.1 (μm or s, only *x* ≥ 0. 5). MLE fails because the complete distribution was not sampled. LSF-CDF(static) method with cutoff of *x_0_* = 0.5 fails because the real cutoff should be at *x_0_* = 0.45 (see Supporting Material S7). All simulations used 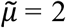 and *n* = 100 (for LSF-PDF a 0.5 bin width was used). Values on the right are mean results of simulations using 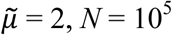, *N*= 10^5^ and *n* = 10^4^.

### Correction for finite filament length

Since each filament has a finite length, some motor proteins are destined to run into the end of their track and detach (Figure 5A). This influences the measurement of run length (or interaction time) and introduces a dependence on the filament length [8]. Therefore, identical motor proteins moving along longer filaments would have a higher observed run length (or interaction time) than motors stepping on short filaments. Here, we present a correction for these so-called ’end-events’ by using the Kaplan-Meier-Estimator [20] in LSF-CDF(free) to adjust the cumulative probability distribution for these censored events. Since it is possible to image the filaments and track single motor proteins, the detachment-positions along the filament can be determined. Any event where a motor protein detaches near the end of a filament (within 1 pixel), is then scored as end-event (the exact procedure to calculate the adjusted cumulative probability distribution can be found in Supporting Material S8). To verify this method, we simulated events of motor proteins landing on filaments with a random length distribution (assuming a Schulz-Distribution [21], equation in Supporting Material S3) and assessed, which motors reach the filament end. These traces are included in the analysis as censored events.

**Figure 5:**
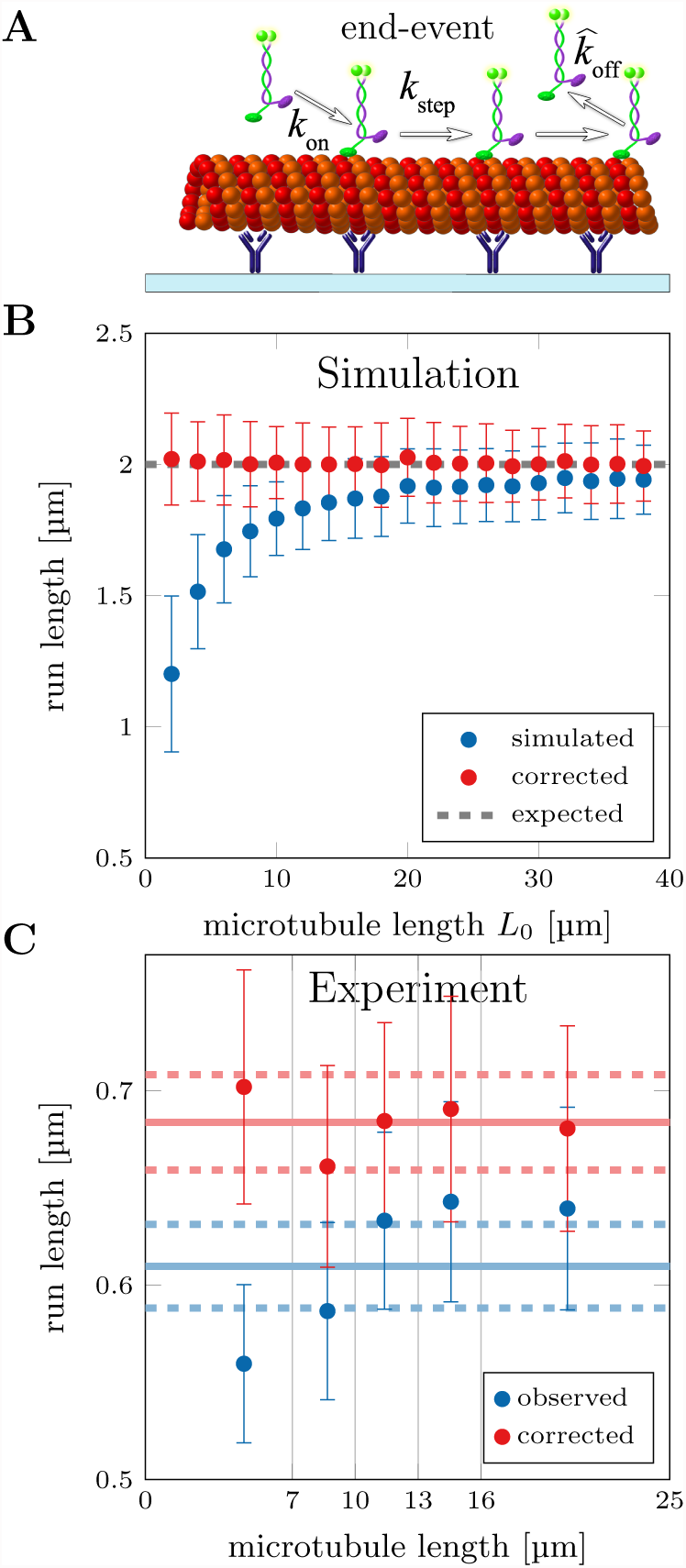
Correction for finite filament length: (A) Schematic depiction of GFP-labeled kinesin-1 motors that reach the end of the microtubule and are forced to detach there (scored as ’end-event’). (B) Simulation of motor proteins landing on 10 filaments picked out of a Schulz distribution with *L*_0_. Some motors reach the end and detach prematurely, which leads to an underestimation of run length. Using the Kaplan-Meier-Estimator (for end-events) a corrected cumulative probability distribution can be calculated and LSFCDF(free) is used to estimate a corrected run length (the same can be applied to the interaction time). (C) Experimental data of kinesin-1 run length, where traces were separated in 5 groups according to the length of their filament (*N* ≈ 1000). Solid line shows mean run length of combined distribution with bootstrapping error (dashed lines, see Methods).

Figure 5B compares the run lengths from our simulated data with and without correction. Note that neglecting the length correction leads to a systematic error that influences the measurement because a certain number of motor proteins will always reach the end even for long filaments. The method was also verified experimentally when we tracked 5208 kinesin-1 motor proteins stepping along MTs (same data as in Figure 2C) in 8 different fields of view (temperature 23.5–24 °C). Figure 5C shows the run lengths grouped according to the length of their MT (combined data set was separated in groups of *N* ≈ 1000). Whereas a dependence of the observed run length on the MT length is seen in the uncorrected data, the correction using the Kaplan-Meier-Estimator yields the same run length for all groups as well as for the combined distribution. The errors can be estimated using bootstrapping, where measurements are randomly selected (with replacement, see Methods), classified for end-events and then evaluated using LSF-CDF(free) with Kaplan-Meier-Estimator. After sufficient repetitions (*n* = 100) the resulting bootstrapping distribution yields the mean run length 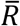 and the statistical error after correction Δ*R* = 2 · *σ_R_*. Therefore, any influence of the Kaplan-Meier-Estimator on the statistical error can be accounted for. An alternative method to verify length correction can be found in the Supporting Material S9.

### Correction for photobleaching

Another experimental limitation is the statistical nature of photobleaching after a fluorophore has emitted a certain number of photons. Photobleaching influences the measurement of the observed interaction time (or run length) and introduces a dependence on the bleaching rate k_bleach_ (see Figure 6A). Even though the lifetime of the fluorophores can be increased by adding antifade solutions [14], the effect of photobleaching can not be eliminated fully in the experiments. Here, we describe the bleaching probability as a combination of one and two fluorophore bleaching (see Eq. 4), because even though dimeric, GFP-labeled motor proteins are always tagged with two fluorophores not all of these fluorophores are active (see Supporting Material S10).

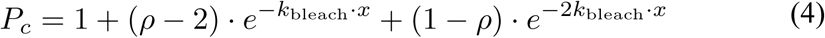

**Figure 6:**
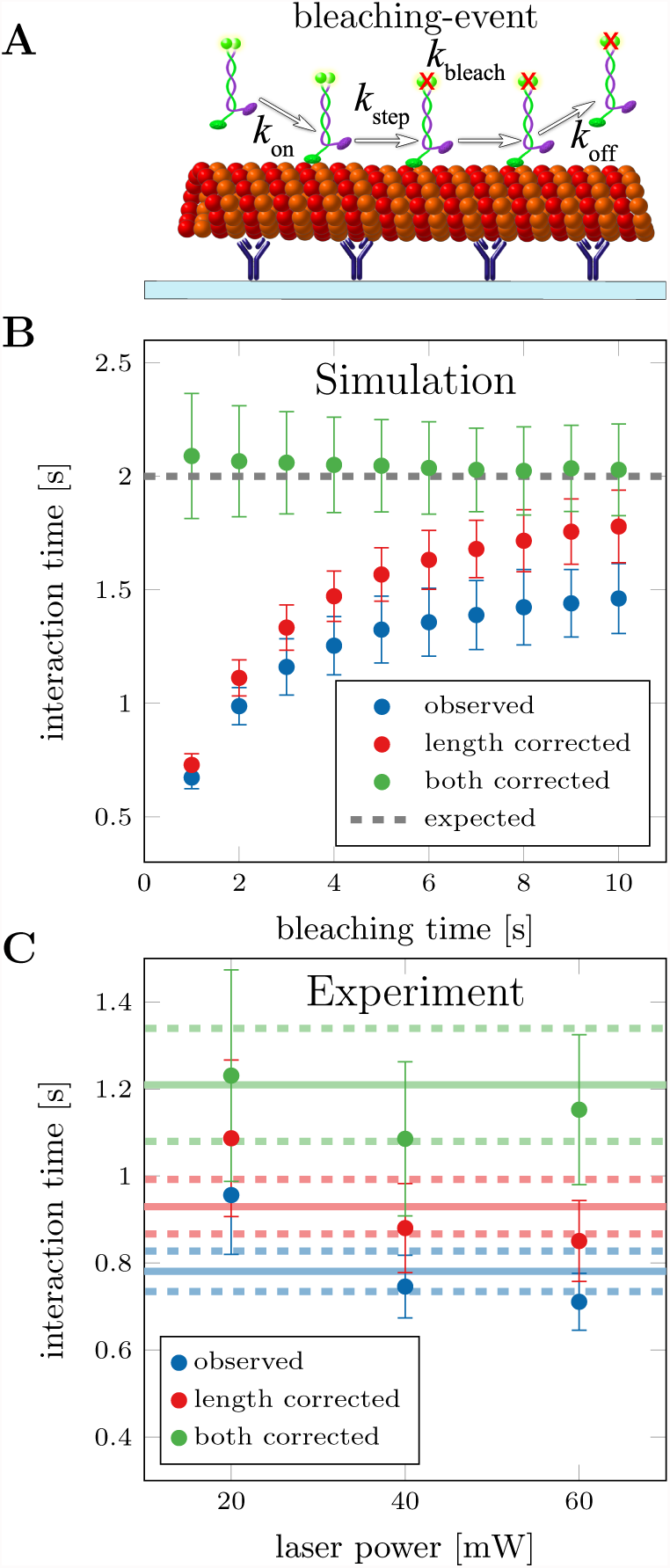
Correction for photobleaching: (A) Schematic depiction of GFP-labeled kinesin-1 motors that photobleach while moving along the microtubule. The observed detachment rate is influenced by photobleaching. (B) Simulations with finite filament lengths (Schulz-Distribution *L*_0_ = 5 μm) and photobleaching of motor proteins with one or two fluorophores (*τ*_bleach_ = 1 – 10 s, *ρ* = 0.5). Distribution of bleaching times (*ρ* and *k*_bleach_) is measured from a different population of simulated motor proteins (e.g. immobilized motors). Simulations used 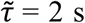, *N* = 1000 and *n* = 100. (C) Experimental data from kinesin-1 at different laser excitation intensities (laser power at controller). The bleaching time distribution was measured with immobilized motor proteins (in the presence of AMP-PNP) in a second flowchannel on the same coverslip (same settings and solutions). Here, bleaching rates *k*_bleach_ and ratio *ρ* were measured for each excitation power: *k*_bleach_ = 0.09 ± 0.02s^-1^ and *ρ* = 0.81 ± 0.06 for 20 mW, *k*_bleach_ = 0.18 ± 0.02 s^-1^ and *ρ* = 0.70 ± 0.06 for 40 mW, *k*_bleach_ = 0.26 ± 0.03s^-1^ and *ρ* = 0.68 ± 0.05 for 60 mW.

In equation 4, the parameter *ρ* denotes the fraction of motors with only one active fluorophore. Because combining the evaluation of detachment and photobleaching is not trivial and the corresponding addition of more parameters into the LSF-CDF leads to unstable solutions, we introduce a different approach by assigning a bleaching probability to each individual motor protein according to its interaction time. Afterwards, the data is analyzed with LSF-CDF(free) several times (*n* = 100) and in each iteration different events will be randomly scored as bleaching-events in agreement with their bleaching probability. Combined with the end-events, these censored events are corrected for by using the Kaplan-Meier-Estimator. This means that the bleaching correction is averaged over many iterations of the run length or interaction time estimation. Additionally, the data can be resampled in every iteration to combine photobleaching correction with the bootstrapping method that now not only yields a corrected measurement, but also the statistical error of the result.

In order to verify the proposed correction, we extended our simulations to include photobleaching with a mixture of one and two fluorophore bleaching *ρ* = 0.5. Figure 6B shows the dependence of the observed and corrected interaction times on the bleaching time. Here, only when correcting for both finite filament length and photobleaching the expected interaction of 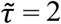 was estimated. Slight deviations from the true value were observed for bleaching times equal to or shorter than the true interaction time, which is due to the limited number measurements that are not censored. We tested the photobleaching correction experimentally by imaging single kineins-1 motor proteins in stepping assays at different laser excitation intensities. Here, for each intensity, the bleaching rate *k*_bleach_ as well as ratio *ρ* was measured in a second flowchannel (on the same coverslip) by immobilizing GFP-labeled kinesin-1 on MTs with AMP-PNP (a non-hydrolysable analog of ATP). The evaluation of the photobleaching using FIESTA is described in the Supporting Material S11. Results in Figure 6C validate the correction for photobleaching, because the dependence of the interaction time on the laser intensity is removed and the estimation of a combined interaction time is possible.

##### Workflow for estimating interaction time and run length

I. Extract the interaction time and run length information for each motor protein (see Supporting Material S2 for instructions on how to track and analyze single fluorescent motor proteins using FIESTA).
II. Score end-events according to their detachment position in relation to the filament end.
III. Estimate bleaching probability from separate channel with immobilized motors (see Supporting Material S11 for photobleaching measurement with FIESTA).
IV. Assign bleaching-events, create cumulative probability distribution (censored end- and bleaching-events) with the Kaplan-Meier-Estimator and use LSF-CDF(free) to estimate interaction time and run length.
V. Bootstrapping: Repeat step IV with different randomly selected traces. Reassign bleaching events randomly in each iteration according to the bleaching probability.

The MATLAB code for evaluation of velocity, bleaching time, interaction time and run length can be found in the Supporting Material S4. Note that censored events also include events where the motor proteins move in or out of the field of view as well as traces that start or end in the first or last imaging frame respectively.

### Discussion

We presented a method to investigate the motility parameters of single, fluorescently-labeled motor proteins in stepping assays, including experimental enhancements, software for tracking and analysis, and precise evaluation of the measurements. For verification, simulations were performed to check whether the methods yield the true results and different methods for evaluation and correction were compared with respect to their systematic and statistical error. The best methods were then used to characterize the temperature dependence of the velocity as well as the influence of the filament length and photobleaching on interaction time and run length experimentally. Furthermore, correction methods are proposed to minimize the influence of the experimental setup on the obtained results. These corrections are shown to work in simulations as well as with experimental data and show the advantage to previously used methods for the evaluation of data from single fluorescent motor proteins.

First, we found that a measured velocity distribution from single motor proteins cannot be described as a normal distribution. When assuming a simple Poisson-Stepper model the velocity of motors with longer interaction times can be estimated more precisely then for short events, which lead to deviations from the normal distribution due to heavier tails. Still, the normal distribution is commonly used when evaluating velocity distributions [3, 4, 5, 6], even though Norris et al. [6] already show clear deviations (Figure S2 in their Supporting Material). Here, we propose using a *t* location-scale (TLS) distribution for estimation of the mean velocity, which fits much better to the simulated as well as the experimentally measured velocity distributions. Compared to the normal distribution this method yields a smaller statistical error. Since it is possible to precisely measure the velocity (Δ*v*/*v* < 0.01 with *N* > 1000) it can be used as a control parameter to verify that the temperature indeed was stable even without the information of the additional temperature sensor in the flow channel.

Secondly, we compared different methods to evaluate exponential distributions and we found that using the least-squares-fitting of the cumulative density function (LSF-CDF) yields the best results. Here, the introduction of a cutoff *x_0_* as a free fit parameter is sufficient to account for missing short events. We note that an adjusted Maximum-Likelihood-Estimation method has recently been published to account for missing short events [22], which was not included in our comparison. Even though it would likely yield better results than the LSF-CDF(free) it is not compatible with our proposed correction of censored events using the Kaplan-Meier-Estimator. These censored events are very specific for stepping assays with fluorescently-labeled motor proteins. Here, detachment of motors at the end of filaments (end-events) and photobleaching of the fluorophores (bleaching-events) distort the results when measuring interaction time and run length for these motor proteins.

Third, we propose specific corrections for finite filament length and photobleaching using the Kaplan-Meier-Estimator with the LSF-CDF(free) method. Since we can track both the fluorescently-labeled motors and the filaments, it is possible to determine end-events, which censor some specific events in a data set. By using the Kaplan-Meier-Estimator we can adjust the cdf for these end-events and thereby correct for the finite filament length. The advantage over previously published methods [8], which use a correction term that includes the average filament length of the experiment, is that the underlying length distribution of the filaments does not influence the evaluation. Therefore, it does not matter if motor proteins are stepping on one particular filament or on a random set of filaments. In addition, variations in filament length between different fields of view do not affect the results. Correction for photobleaching is not as trivial as the filament length correction. Previously, corrections for photobleaching assumed the simple relation *k*_observed_ = *k*_off_ + *k*_bleach_ [9, 23], with *k*_observed_ describing the observed detachment rate being the superposition of the real detachment rate *k*_off_ and the bleaching rate *k*_bleach._ There, by either estimating the bleaching rate [9] or extrapolating *k*_observed_ using different laser intensities [23], a corrected interaction time could be calculated. Unfortunately, these methods fail when the molecules of interest are labeled with more than one fluorophore as is the case with dimeric motor proteins where a GFP is expressed on each monomeric motor unit. Most importantly, not all fluorophores are active, which leads to a mixture of observed motor proteins with either one or two fluorophores. Therefore, we propose to measure the bleaching time as well as the ratio of one to two fluorophore bleaching, which enables us to calculate a bleaching probability for a specific experiment. Over many iterations, we can now randomly assign bleaching-events according to the interaction time of the motor proteins. By including them in the censored events, our correction for both, photobleaching and finite filament length, is achieved. Consequently, our simulations show slight deviations from the expected true values only for bleaching times in the range of - or shorter than - the interaction time.

How well does our proposed method perform when (i) the interaction time of the motor is significantly longer than the bleaching time of the fluorophores, (ii) the run lengths of the motors are significantly larger than the filament lengths, and (iii) the motors exhibit frequent switching between movement and stalling? With regard to the limitations originating from photobleaching, adjustment of the imaging conditions (e.g. exposure time and light intensity, such that the bleaching rates are lower than the motor detachment rates) are effective means to reduce the systematic error. Furthermore, novel photo-stable fluorescent proteins as well as functionalized fluorophores (e.g. GFP-boosters based on nanobodies) will reduce the influence of photobleaching. In any case, comparing the motility parameters obtained under different imaging conditions can directly show if photobleaching influenced the results. With regard to investigating highly-processive motor proteins, which reach the filament ends in most of the cases, the only solution is to use longer MTs. Tweaking the protocols for MT polymerization (e.g. by extending the growths times in conjunction with lowering the tubulin concentration) allows for the generation of MTs with lengths above 50 μm, sufficient to reliably determine the run lengths in any cellular context. Moreover, discarding all motility events from filaments with lengths below a threshold is possible, because our proposed correction algorithms do not require a certain MT length distribution. In general, the more motor traces are censored the less accurate the precision will be. With respect to super-processive motility (such as Kinesin-3 [24] and Kinesin-8 [25]) we note that the run length is not a suitable motility parameter and rather other measures (such as the motor’s end-reach probability in dependence of the landing position) should be applied. With regard to motors which exhibit frequent stalling, our corrections for finite filament length and photobleaching do stay valid (Kaplan-Meier-Estimator can still be used), but the motors cannot be described by simple Poisson steppers anymore. Therefore, the underlying models will have to be adjusted accordingly, e.g. by using double-exponential functions for interaction time and run length.

We conclude, in order to precisely characterize the stepping of motor proteins on their filaments the following steps are necessary: (i) The temperature should be stable throughout the experiments in order to combine and evaluate many traces at the same condition in one data set. Because even small changes in temperature influence the motility paramaters significantly, it is essential to measure the assay temperature precisely. Measuring and specifying the room temperature is unsuited for this purpose, as the actual temperature in the flow channel can be up to 3 K higher due to microscope-internal heat sources, such as electrical components and light sources. Hence, any results given for velocity, run length and interaction time should include the temperature in the flow channel within ±1 K. Furthermore, in order to investigate different motor or filament populations, we recommend to incorporate them in the same flow channel or at least on the same coverslip in order to minimize any temperature differences in the experiments. (ii) Corrections for finite filament length and photobleaching need to be included, otherwise both interaction time and run length are underestimated (e.g. see Zimmermann et al. [26]). Here, the systematic errors can be on the same order as the statistical error and therefore careful consideration of the evaluation method is essential. For that reason, we provide an extensive description of the evaluation method, including the MATLAB code, to efficiently measure a sufficient number of motor proteins (to reduce the statistical error) and to address limitations in the design of the experimental assay (to remove the systematic error). (iii) The motility parameters velocity, interaction time and run length should always be estimated when comparing different data sets. Motor proteins could have the same run length, but different velocities (*k*_step_) and interaction times (*k*_off_), so only comparing one motility parameter might result in the misconception that even if the experiment yields the same result for differenct motors the underlying motility mechanism could still be different. This allows for a better statistical comparison of motor proteins influenced by external factors e.g. ionic strength, ATP concentration, nucleotide state of the filaments or post-translational modifications of the filaments. Furthermore, comparison of different motor proteins as well as motor populations, e.g. structural differences or binding of regulatory proteins, will then become possible. We believe, the methodology developed in our work will provide a reliable framework for the evaluation of a wide range of experiments with single fluorescently-labeled motor proteins.

## Author Contributions

Conceived and designed the experiments: FR SD. Performed the experiments: FR LK. Analyzed the data: FR LK. Contributed reagents/materials/analysis tools: FR. Wrote the paper: FR SD.

## Acknowledgments

We thank Friedrich Schwarz for help with the design and the mechanical workshop at the MPICBG for manufacturing the cooling/heating ring for the objective, Corina Bräuer for technical support, Aniruddha Mitra, Jens Ehrig, Rahul Grover and Georg Krainer for comments on the manuscript and the whole Diez-Lab for fruitful discussions. We acknowledge financial support from the European Research Council (Starting Grant 242933), the DFG through the Center for Advancing Electronics Dresden (cfaed) and the Technische Universität Dresden.

